# Comparison of urine proteome between obese people and normal weight people

**DOI:** 10.1101/2024.01.06.574495

**Authors:** Haitong Wang, Lilong Wei, Yun Zhou, Yongtong Cao, Youhe Gao

## Abstract

**Objective:** To explore whether urine proteome can reflect the difference between obese and normal weight people.

**Methods:** Urine samples from obese and normal weight people were collected and identified by non-label quantitative proteomics using high performance liquid chromatography tandem mass spectrometry (LC-MS/MS). The difference proteins of urine proteome between obese and normal weight people were screened for protein function and biological pathway analysis. The urine proteome of obese individuals was compared with that of normal weight people, and the common differential proteins were counted to analyze the protein function and biological pathways. Reported biomarkers of obesity were searched in the urine proteome of obese individuals.

**Results:** 38 different proteins can be identified in the urine proteome of obese people compared with normal weight people, some of which have been reported to be related to metabolism and obesity, and the biological processes of differential proteins are also related to metabolism and other processes. 8 common differential proteins in the urine proteome of obese individuals and normal weight people, among which some proteins have been reported to be related to metabolism and obesity, and the biological processes of differential proteins are also related to metabolism and other processes. Among the differential proteins in the urine proteome of obese individuals compared with the normal weight people, the reported obesity biomarkers can be matched.

**Conclusions:** The urine proteome can distinguish the obese people from the normal weight people, and the differential proteins in the urine proteome have key proteins that are known to be related to obesity and metabolism, and the biological processes of differential proteins also related biological processes such as nutrition and metabolism. Urine proteome has the potential to explore the pathogenesis of obesity and provide personalized treatment.

## 1 Introduction

The global obesity epidemic is a major public health problem today. Obesity increases the risk of many chronic diseases, such as type 2 diabetes, coronary heart disease and cancer. The pathogenic factors of obesity are complex, result from the interaction of genetic, nutritional and metabolic factors. Although there is no clear consensus on the terms of classification and characterization of obesity, it can be roughly divided into the following four phenotypes: (1) normal weight obesity (NWO); (2) metabolically obese normal weight (MONW); (3) metabolically healthy obese (MHO); (4) metabolically unhealthy obese (MUO); (5) sarcopenic obesity (SO)[1 2].

At present, there is no significant biomarker used to distinguish obesity and each obesity subtype. The Body Mass Index (BMI) used to classify obesity is calculated by dividing weight (kg) by the square of height (m). According to the Chinese BMI classification, Underweight was defined as BMI < 18.5 kg/m2, normal weight as 18.5 kg/m2 ≤ BMI < 24 kg/m2, overweight as 24 kg/m2 ≤ BMI < 28 kg/m2, obesity as BMI ≥ 28 kg/m2. Is an imperfect measure of abnormal body fat [1 3 4].

More refined techniques, such as magnetic resonance imaging, are used to assess body fat distribution to better diagnose subtypes of obesity. However, these tests are not easy to perform in routine clinical Settings, and cut-off values have not been established [1]. However, a simpler approach to rapidly identify specific biomarkers to characterize the causes of obesity and provide targets in personalized treatment still needs further research. Urine is produced by blood filtration through the kidney to exclude metabolic waste from the body. Since it does not belong to the internal environment of the body and is not controlled by homeostatic regulation mechanism, it can retain very small physiological changes of the body [5]. Studies have shown that urine proteome can detect biomarkers of diseases, such as diabetes [6], Alzheimer’s disease [7], depression [8], autism [9]; Biomarkers of urine proteome can also classify diseases, such as predicting chronic kidney disease [10] and distinguishing benign and malignant ovarian cancer [11]. However, urine proteomics has not yet been used to find biomarkers for the causes of obesity and personalized treatment. This study is the first to propose a urine proteomic analysis of obese and normal weight people to explore whether urine can reflect the causes of obesity, provide potential drug targets, and assist in personalized treatment.

## 2 Experimental method

### 2.1 Sample collection

A total of 19 urine samples were collected from the China-Japan Friendship Hospital in Beijing. According to the Chinese reference standard of BMI, 18.5 kg/m^2^ < BMI < 23.9 kg/m^2^ is normal weight; BMI > 28 kg/m^2^ is obese. There were 10 samples in the obese group, with an average BMI of 35.79 kg/m^2^.There were 9 samples in the normal weight group, with an average BMI of 22.76 kg/m^2^. This experiment is based on the reuse of discarded samples in the laboratory, and the process does not involve any identity information of patients. Ethics review number: 2023-KY-126.The BMI of the subjects is shown in Table 1:

**Table 1.** BMI of obese group and normal weight group.

| | ID | Age | BMI ($\text{kg/m}^2$) |
| --- | --- | --- | --- |
| Normal weight group | 1 | 32 | 21 |
| Normal weight group | 2 | 31 | 21 |
| Normal weight group | 3 | 34 | 22 |
| Normal weight | 4 | 33 | 23 |

| group |  |  |  |
| --- | --- | --- | --- |
| Normal weight group | 5 | 33 | 23. |
| Normal weight group | 6 | 34 | 23 |
| Normal weight group | 7 | 33 | 24 |
| Normal weight group | 8 | 35 | 24 |
| Normal weight group | 9 | 27 | 24 |
| Obese group | 11 | 33 | 30 |
| Obese group | 12 | 33 | 30 |
| Obese group | 13 | 30 | 30 |
| Obese group | 14 | 29 | 31 |
| Obese group | 15 | 31 | 32 |
| Obese group | 16 | 33 | 34 |
| Obese group | 17 | 28 | 37 |
| Obese group | 18 | 29 | 41 |
| Obese group | 19 | 19 | 44 |
| Obese group | 20 | 19 | 49 |

### 2.2 Urine sample processing

Urine protein extraction: Urine samples were removed from the refrigerator at -80 °C and thawed at 4°C. Centrifuge at 4°C, 12000×g for 30 min, take 6 mL of supernatant into 50 mL centrifugal tube, add 20 mM Dithiothreitol (DTT, Sigma), swirl and mix, heat in a water bath at 37°C for 60 min, and cool to room temperature. Add 50 mM Iodoacetamide (IAA, Sigma), swirl and mix, and react at room temperature for 40 min away from light. Add triploid volume of pre-cooled anhydrous ethanol, gently mix upside down, and precipitated at -20°C for 24 h. The mixture precipitated 24 h was centrifuged at 4°C, 12000×g for 30 min, the supernatant was discarded, and the ethanol was volatilized and dried. The protein precipitate was suspended in the lysate (containing 8 mol/L urea, 2 mol/L thiourea, 25 mmol/L dithiothreitol, 50 mmol/L Tris). Centrifuge at 4°C at 12000×g for 30 min, supernatant was taken into a new 1.5 mL centrifuge tube to obtain urine protein. The protein concentration was determined by Bradford method.

Enzyme digestion of urine protein: 100 μg urine protein sample was taken into 1.5 mL centrifuge tube, and 25 mmol/L NH_4_HCO_3_ solution was added to make the total volume 200 μL. Take a 10 kDa ultrafiltration tube (Pall, Port Washington, NY, USA) and add 200 μL UA solution (8 mol/L urea, 0.1 mol/L Tris-HCl, pH 8.5) to the filter membrane to wash the filter membrane. Centrifuge at 18°C, 14000×g for 5 min, discard the lower filtrate, and repeat once; Urine protein samples were added to the filter membrane, centrifuged at 18°C, 14000×g for 30 min, and the lower filtrate was discarded, leaving urine protein on the filter membrane. Urine protein was washed with 200 μL UA solution into the filter, centrifuged at 18°C, 14000×g for 30 min, and repeated twice. Urine protein was washed with 25 mmol/L NH_4_HCO_3_ solution into the filter membrane, centrifuged at 18°C, 14000×g for 30 min, and repeated twice. Trypsin Gold (Promega, Fitchburg, WI, USA) was added at the ratio of 1:50 for enzyme digestion, and heat in a water bath at 37°C for 16 h. After digestion, the filtrate was collected by centrifugation at 4°C, 13000×g for 30 min. The filtrate was a polypeptide mixture. The polypeptide mixture was desalted by HLB solid phase extraction column (Waters, Milford, MA), lyophilized with a vacuum dryer, and stored at -20°C.

### 2.3 LC-MS/MS tandem mass spectrometry analysis

Lyophilized polypeptide mixture with 0.1% formic acid and quantified with BCA kit. The peptide concentration was diluted to 0.5 μg/μL. Each sample was mixed with 6 μL and separated by Thermo Fisher Scientific (high pH reversed phase peptide separation kit). Centrifuge and collect 10 Fractions, freeze dried by vacuum dryer and redissolve in 0.1% formic acid. An iRT reagent (Biognosys, Switzerland) was added to 10 effluents and all individual samples with a sample: iRT volume ratio of 10:1 to calibrate the retention time of the extracted peptide peaks.

10 Fractions were separated using the EASY-nLC1200 chromatography system (Thermo Fisher Scientific, USA). The isolated peptides were analyzed by Orbitrap Fusion Lumos Tribrid mass spectrometer (Thermo Fisher Scientific, USA) in Data Dependent Acquisition (DDA) mode. Generate 10 raw files and import them into Proteome Discoverer software for database construction analysis using Swiss-iRT and Uniprot-Human databases (version 2.0, Thermo Scientific). The DIA method is established by setting 39 variable Windows of the Data Independent Acquisition (DIA) model of a single sample according to the results of the database construction. A single sample of 1 μg peptide was isolated using EASY-nLC1200 chromatography (Thermo Fisher Scientific, USA). The separated peptides were analyzed by Orbitrap Fusion Lumos Tribrid mass spectrometer (Thermo Fisher Scientific, USA) in DIA mode, the newly established DIA method was used to collect DIA data and generate raw files.

### 2.4 Quantitative analysis of Label-free DIA

The raw file of a single sample collected in DIA mode was imported into Spectronaut Pulsar(Biognosys AG, Switzerland) for analysis. The abundances of peptide segments were calculated by summing the peak areas of each fragment ion in MS2. Protein abundance is calculated by adding the abundances of the respective peptide segments.

### 2.5 Data analysis

The technique was repeated 3 times for each sample, and the average value was taken for statistical analysis. In this experiment, obese group and normal weight group were compared and screened for different proteins. The screening conditions for differential proteins were: Fold change (FC) ≥ 2 or ≤ 0.5, P < 0.01 for double-tailed unpaired T-test analysis; At the same time, a one-to-many comparative analysis was performed in this experiment. A single sample in the obese group was compared with 9 samples in the normal weight group, and differential proteins were screened. The screening conditions for differential proteins were: FC ≥ 1.5 or ≤ 0.67, P value < 0.05 for double-tailed unpaired T-test analysis. The common differential proteins shared by 10 samples in the obese group were counted. Differential proteins were analyzed on the website of Uniprot (https://www.uniprot.org/) and DAVID database (https://david.ncifcrf.gov/), related literatures were searched in Pubmed database (https://pubmed.ncbi.nlm.nih.gov) for functional analysis.

## 3 Experimental results and discussion

### 3.1 Comparison of urine proteome between obese group and normal weight group

#### 3.1.1 Differential proteins

The urine proteins of the obese group and the normal weight group were compared, and the screening conditions for Differential proteins were FC ≥ 2 or ≤ 0.5, and double-tail unpaired T-test P < 0.01. The results showed that, compared with the normal weight group, 38 differential proteins could be identified in the obese group. The differential proteins were arranged in the order of FC from small to large, and the results were retrieved by Uniprot, as shown in Table 2.

**Table 2.**
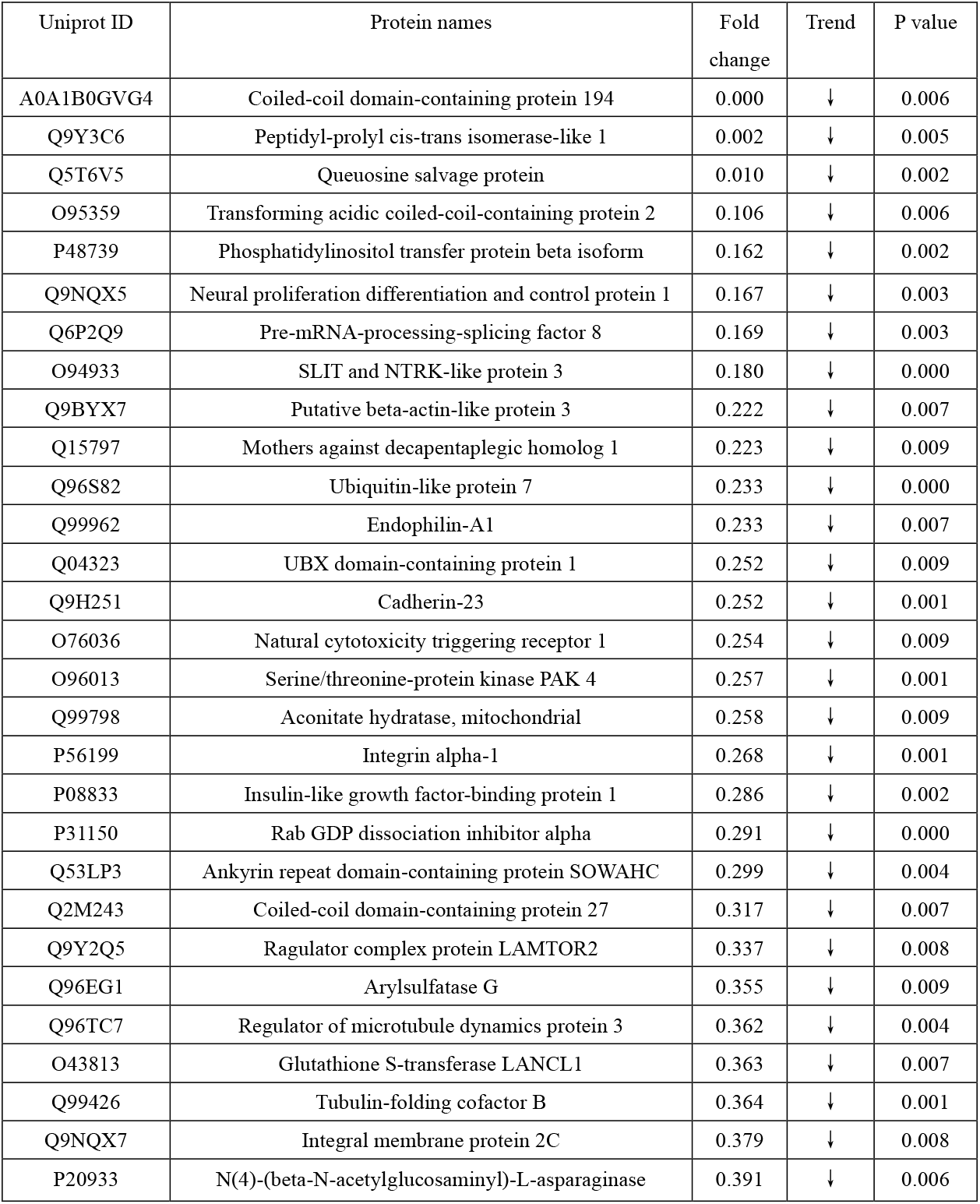
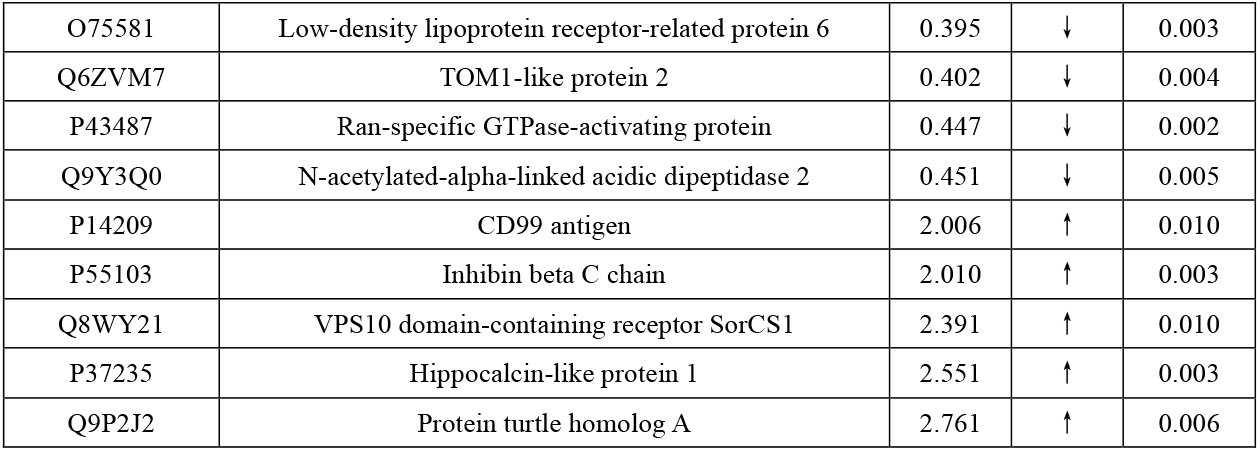
Urine proteomic differences between obese group and normal weight group.

| Uniprot ID | Protein names | Fold change | Trend | P value |
| --- | --- | --- | --- | --- |
| A0A1B0GVG4 | Coiled-coil domain-containing protein 194 | 0.000 | ↓ | 0.006 |
| Q9Y3C6 | Peptidyl-prolyl cis-trans isomerase-like 1 | 0.002 | ↓ | 0.005 |
| Q5T6V5 | Queuosine salvage protein | 0.010 | ↓ | 0.002 |
| O95359 | Transforming acidic coiled-coil-containing protein 2 | 0.106 | ↓ | 0.006 |
| P48739 | Phosphatidylinositol transfer protein beta isoform | 0.162 | ↓ | 0.002 |
| Q9NQX5 | Neural proliferation differentiation and control protein 1 | 0.167 | ↓ | 0.003 |
| Q6P2Q9 | Pre-mRNA-processing-splicing factor 8 | 0.169 | ↓ | 0.003 |
| O94933 | SLIT and NTRK-like protein 3 | 0.180 | ↓ | 0.000 |
| Q9BYX7 | Putative beta-actin-like protein 3 | 0.222 | ↓ | 0.007 |
| Q15797 | Mothers against decapentaplegic homolog 1 | 0.223 | ↓ | 0.009 |
| Q96S82 | Ubiquitin-like protein 7 | 0.233 | ↓ | 0.000 |
| Q99962 | Endophilin-A1 | 0.233 | ↓ | 0.007 |
| Q04323 | UBX domain-containing protein 1 | 0.252 | ↓ | 0.009 |
| Q9H251 | Cadherin-23 | 0.252 | ↓ | 0.001 |
| O76036 | Natural cytotoxicity triggering receptor 1 | 0.254 | ↓ | 0.009 |
| O96013 | Serine/threonine-protein kinase PAK 4 | 0.257 | ↓ | 0.001 |
| Q99798 | Aconitate hydratase, mitochondrial | 0.258 | ↓ | 0.009 |
| P56199 | Integrin alpha-1 | 0.268 | ↓ | 0.001 |
| P08833 | Insulin-like growth factor-binding protein 1 | 0.286 | ↓ | 0.002 |
| P31150 | Rab GDP dissociation inhibitor alpha | 0.291 | ↓ | 0.000 |
| Q53LP3 | Ankyrin repeat domain-containing protein SOWAHC | 0.299 | ↓ | 0.004 |
| Q2M243 | Coiled-coil domain-containing protein 27 | 0.317 | ↓ | 0.007 |
| Q9Y2Q5 | Ragulator complex protein LAMTOR2 | 0.337 | ↓ | 0.008 |
| Q96EG1 | Arylsulfatase G | 0.355 | ↓ | 0.009 |
| Q96TC7 | Regulator of microtubule dynamics protein 3 | 0.362 | ↓ | 0.004 |
| O43813 | Glutathione S-transferase LANCL1 | 0.363 | ↓ | 0.007 |
| Q99426 | Tubulin-folding cofactor B | 0.364 | ↓ | 0.001 |
| Q9NQX7 | Integral membrane protein 2C | 0.379 | ↓ | 0.008 |
| P20933 | N(4)-(beta-N-acetylglucosaminy)-L-asparaginase | 0.391 | ↓ | 0.006 |
| O75581 | Low-density lipoprotein receptor-related protein 6 | 0.395 | ↓ | 0.003 |
| Q6ZVM7 | TOM1-like protein 2 | 0.402 | ↓ | 0.004 |
| P43487 | Ran-specific GTPase-activating protein | 0.447 | ↓ | 0.002 |
| Q9Y3Q0 | N-acetylated-alpha-linked acidic dipeptidase 2 | 0.451 | ↓ | 0.005 |
| P14209 | CD99 antigen | 2.006 | ↑ | 0.010 |
| P55103 | Inhibin beta C chain | 2.010 | ↑ | 0.003 |
| Q8WY21 | VPS10 domain-containing receptor SorCS1 | 2.391 | ↑ | 0.010 |
| P37235 | Hippocalcin-like protein 1 | 2.551 | ↑ | 0.003 |
| Q9P2J2 | Protein turtle homolog A | 2.761 | ↑ | 0.006 |

#### 3.1.2 Functional analysis of differential proteins

The 38 identified differential proteins were searched through PubMed database.

Among them, some of the most significant differential proteins have not been reported to be associated with obesity, diabetes and other diseases. However, among the differential proteins, Insulin-like growth factor-binding protein 1 (IGFBP-1), FC= 0.286, P = 0.002, that is, the obese group was down-regulated more than 3 times compared with the normal weight group. A large number of studies have shown that this protein is negatively correlated with BMI, waist-to-hip ratio, and fasting insulin level [12 13].

IGFBP-1 levels are dynamically regulated by insulin.When postprandial Insulin is secreted in large amounts, it will inhibit the upstream promoter of IGFBP-1, inhibit its expression, and the level of IGFBP-1 will decrease rapidly, thus increasing the biological activity of insulin-like growth factor 1 and enhancing its insulin-like effect [14, 15, 16].

It has been demonstrated in different populations such as Europeans and Pakistanis [17], Asian Indians [18], healthy young people [19], adults over 65 years old [20], obese menopausal women [21], patients with type 1 diabetes [22] and prepubertal children [12, 23], IGFBP 1 concentration were positively correlated with insulin sensitivity. Therefore, IGFBP-1 is considered as a potential marker of insulin sensitivity [24]. Through a study of 615 individuals, it was determined that IGFBP-1 concentration and its interaction with IGF-1 are important determinants of glucose intolerance or the development of diabetes [25]. A study of 355 Swedish men demonstrated that low fasting IGFBP-1 concentration predicted the development of abnormal glucose regulation, some of whom had a 40-fold increased risk of diabetes [26]. Another study followed 782 individuals for 17 years and found that low expression of IGFBP-1 predicted the development of type 2 diabetes [27]. A study of 240 women over 8 years also showed that IGFBP-1 was associated with an increased risk of diabetes [28].

In addition, other differential proteins have been reported to be associated with metabolism or obesity. Aconitate hydratase is involved in tricarboxylic acid cycle and carbohydrate metabolism, catalyzes the isomerization of citrate into isocitrate through cisaconite acid, and can control adipogenesis by mediating the production of cellular ATP [29]. Inhibin and activin can inhibit and activate pituitary secretion of follicle-stimulating hormone, respectively. Inhibin beta is involved in the regulation of many functions, such as hypothalamic and pituitary hormone secretion, gonadal hormone secretion, germ cell development and maturation, red blood cell differentiation, insulin secretion, nerve cell survival, embryonic axial development or bone growth, and BMI is an important independent predictor of its level [30].

Some differential proteins between the obese group and the normal weight group have been reported to be associated with obesity or metabolism; while some differential proteins with more significant changes than these proteins have not been reported to be associated with obesity or metabolism. It is worthwhile to further investigate the function of significant changes in differential proteins in obesity and metabolism in order to search for biomarkers of obesity or potential drug targets.

#### 3.1.3 Analysis of biological processes of differential proteins

Biological pathways were analysed for differential proteins between the obese and normal weight groups using the DAVID database. A total of 24 biological processes with p < 0.01 were enriched and the results are shown in Figure 1.

**Fig. 1.**
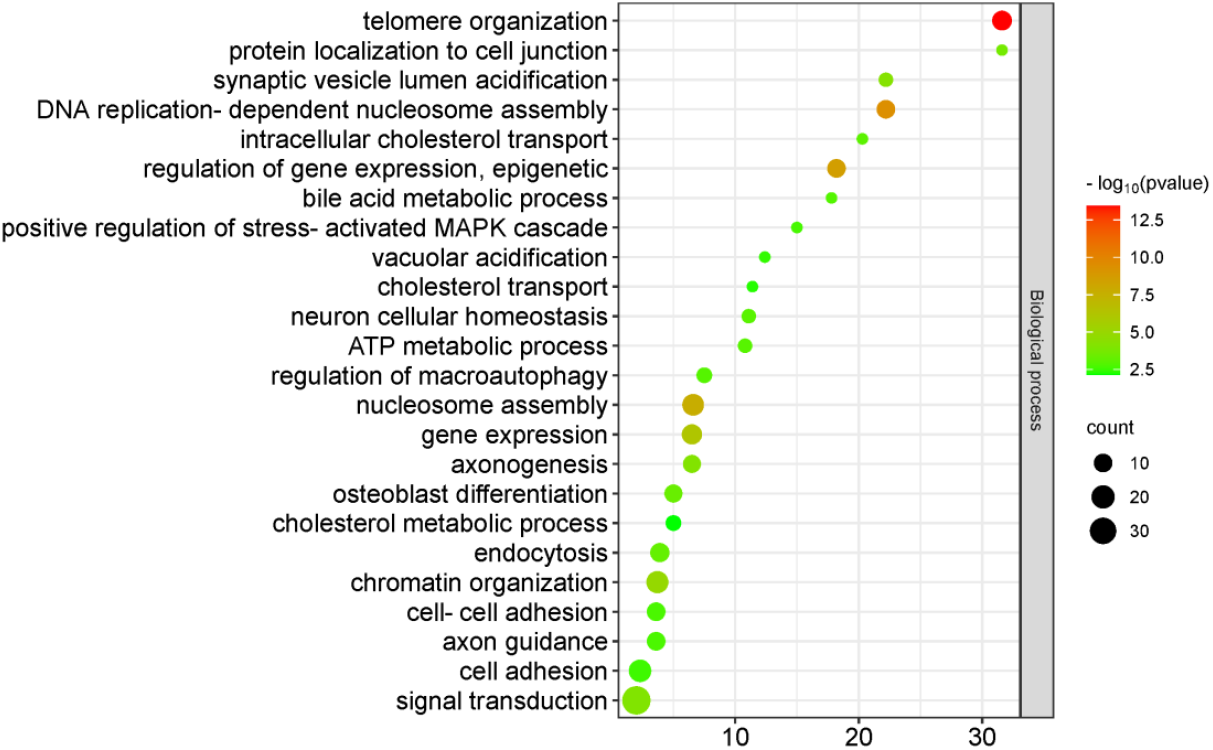
Biological process of differential proteins in urine proteome between obesity group and normal weight group.

These include a number of metabolism-related biological processes, such as intracellular cholesterol transport, ATP metabolic process, bile acid metabolic process, cholesterol transport and cholesterol metabolic process. However, there are other biological processes related to gene expression and the nervous system, which have smaller P-values relative to metabolism-related biological processes. Among them telomere organisation may be related to obesity, and it has been shown that telomere length is a strong marker of biological aging and increased telomere wear has been noted in obese adults [31]. The rest of the biological processes have not been found to be associated with obesity or BMI at this time, which provides new research directions and ideas for understanding the causes of obesity.

### 3.2 Comparison of urine proteome between individual individuals in obese group and normal weight group

#### 3.2.1 Individual shared common differential proteins in the obese group

The urine proteome of a single individual in the obese group was compared with the normal weight group to screen for differential proteins.The conditioned were FC ≥ 1.5 or ≤ 0.67, and a two-tailed unpaired t-test of P < 0.05 was performed to statistically identify differential proteins shared by individuals in the obese group. The results showed that 8 shared common differential proteins were identified, as shown in Table 3. Notably, all differential proteins of single individuals in the obese group were decreased relative to the normal weight group, and except for Peptidyl-prolyl cis-trans isomerase-like 1 and Queuosine 5’-phosphate N-glycosylase/hydrolase, the remaining 6 differential proteins were expressed in the normal weight group, while none of them were expressed in individual individuals of the obese group.

**Table 3.** The shared common differential proteins of urine proteome of individual individuals from the obese group compared with the normal weight group.

| Uniprot ID | Protein names | Trend |
| --- | --- | --- |
| Q9Y5F0 | Protocadherin beta-13 | ↓ |
| Q8IY92 | Structure-specific endonuclease subunit SLX4 | ↓ |
| Q96AA8 | Janus kinase and microtubule-interacting protein 2 | ↓ |
| Q9Y3C6 | Peptidyl-prolyl cis-trans isomerase-like 1 | ↓ |
| A0A1B0GVG4 | Coiled-coil domain-containing protein 194 | ↓ |
| P47914 | Large ribosomal subunit protein eL29 | ↓ |
| Q5T6V5 | Queuosine 5'-phosphate N-glycosylase/hydrolase | ↓ |
| P51151 | Ras-related protein Rab-9A | ↓ |

#### 3.2.2 Shared common differential protein function analysis

Although the changes of shared common differential proteins in the obese group relative to the normal weight group were significant and the trend was uniform, most of these proteins have not been reported to be associated with diseases such as obesity and diabetes. Especially Coiled-coil domain-containing protein 194, the function of this protein and the biological process it is involved in cannot be found on the Uniprot website. Even so, some of the shared common differential proteins have been associated with obesity, e.g. Protocadherin beta has calcium-binding capacity and may be involved in the establishment and maintenance of specific neurons in the brain; the Protocadherin beta gene is expressed in the hypothalamus and is capable of rare mutations that have biological effects. However, in a study involving 30 extremely obese white adult subjects (mean BMI = 51.1 kg/m2), the detection of peripheral blood white blood cells showed that this rare mutation was significantly enriched, while this phenomenon was not found in the normal weight BMI population [32]. Rare mutations in the same gene were significantly enriched in another study involving obese Korean children [33]. When detected by urinary proteomic, the protein was detectable in the obese group with a mean BMI= 35.79 kg/m^2^ which was significantly different from the normal weight group, side by side illustrating the sensitivity of urinary proteomics as well as demonstrating the potential of urinary proteomics in finding early markers of obesity. The extremely obese population was also enriched for mutations in the Olfactory receptor gene, and variants in both genes have a synergistic effect in extreme obesity [34].

These differential proteins showed significant changes and extremely identical trends in the obese group, but the studies on all of these proteins are still relatively scarce, therefore, these differential proteins deserve further investigation of their functions in obesity and metabolism, period, in order to search for biomarkers or potential drug targets for obesity.

#### 3.2.3 Shared common differential protein function analysis

Biological pathways were analysed using the DAVID database for differential proteins shared by 8-10 individuals in the obese group. A total of 15 biological processes with P < 0.05 were enriched and the results are shown in Figure 2.

**Fig. 2.**
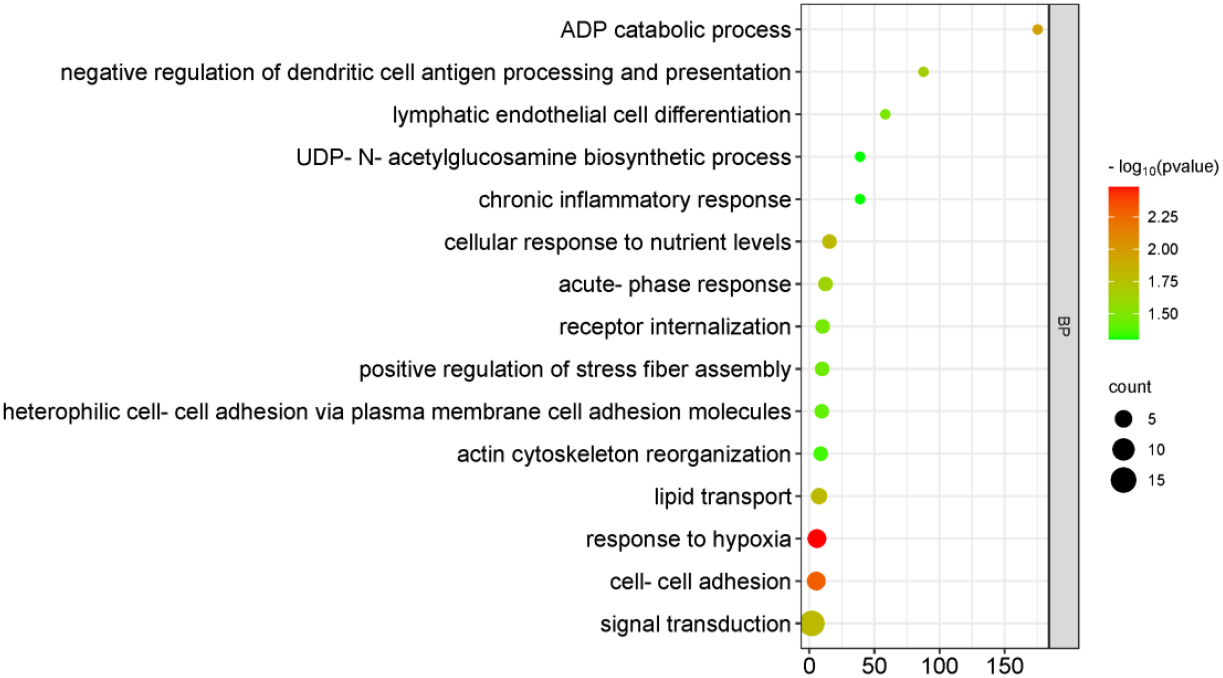
Biological processes of differential protein enrichment shared by 8-10 individuals in obese group.

These biological processes related to nutrition and metabolism, such as: response to hypoxia, which has been shown to exacerbate adipose tissue dysfunction and stimulate the secretion of inflammatory molecules, which can lead to obesity [35], ADP catabolic process, cellular response to nutrient levels, lipid transport and cholesterol metabolic process, UDP-N-acetylglucosamine biosynthetic process, adipose tissue UDP-N-acetylglucosamine was significantly positively correlated with BMI that inhibits UDP-N-acetylglucosamine biosynthesis, leading to reduced glucose-stimulated leptin release in cultured adipocytes [36]. Some studies have shown that dysregulation of glucose metabolism associated with obesity, diabetes or cancer correlates with increased levels of the enzyme in this process. But there are other biological processes involving gene expression and neurological correlations that have smaller P-values relative to these biological processes. One of them, telomere organisation, may be associated with obesity, and it has been shown that telomere length is a strong marker of biological aging and increased telomere wear has been noted in obese adults [30]. The rest of the biological processes have not been found to be associated with obesity or BMI at this time, which provides new research directions and ideas for understanding the causes of obesity.

### 3.3 Personality analysis of individuals in the obese group

A large number of obesity biomarkers have been identified in existing studies [1 37 38]. These markers were searched for in the urine proteome differential proteins of a single individual in the obese group versus the normal weight group. Many markers can be used to differentially react through the urinary proteome, and the markers appear differently in different individuals, as shown in Table 4. This suggests that the urine proteome has the potential to assist in the development of personalised treatment plans for obesity.

**Table 4.**
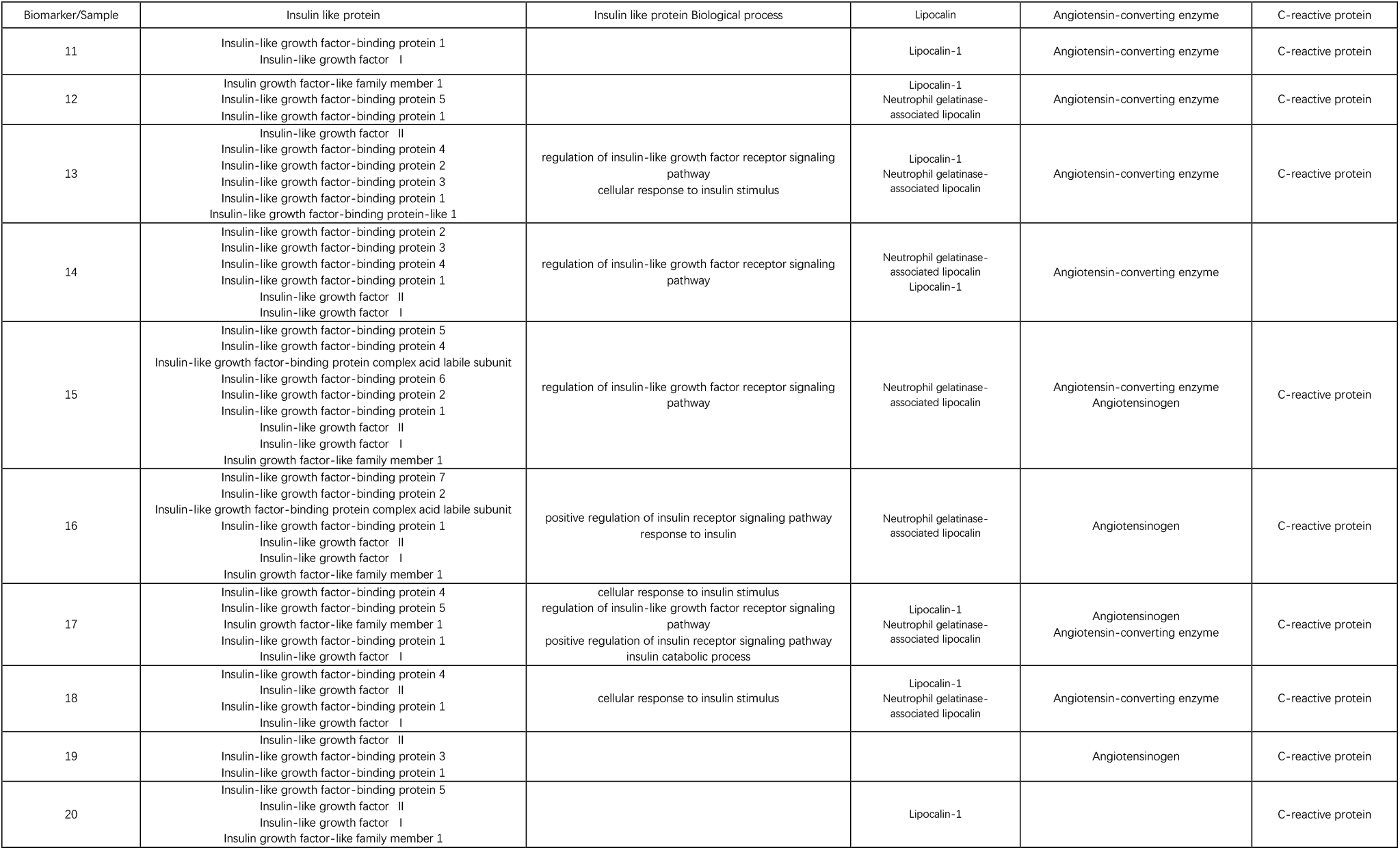
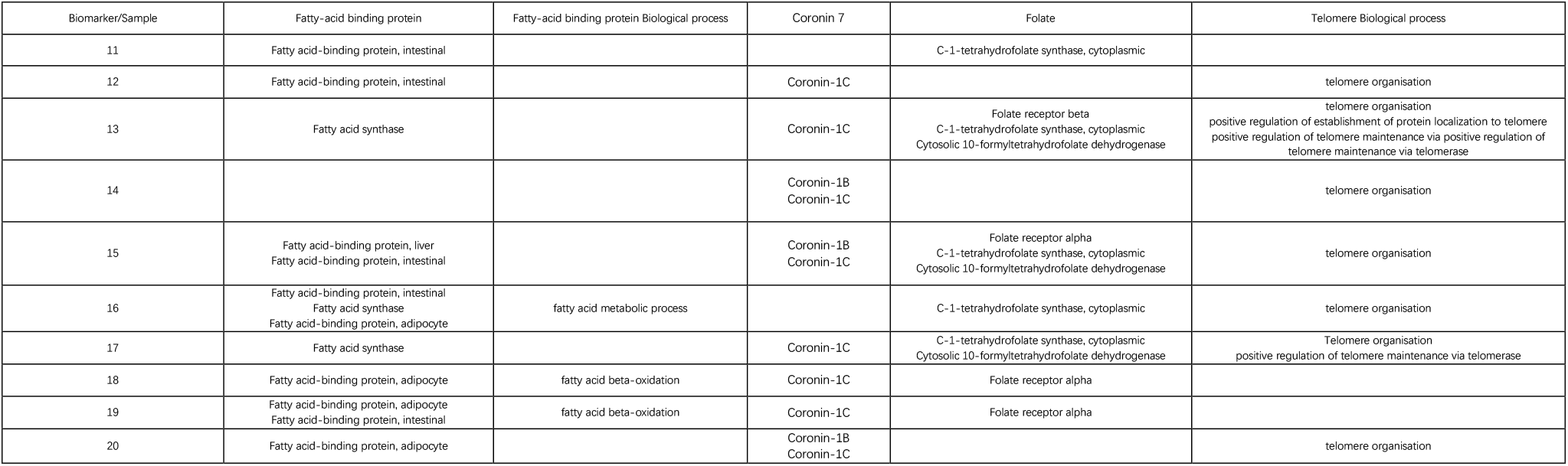
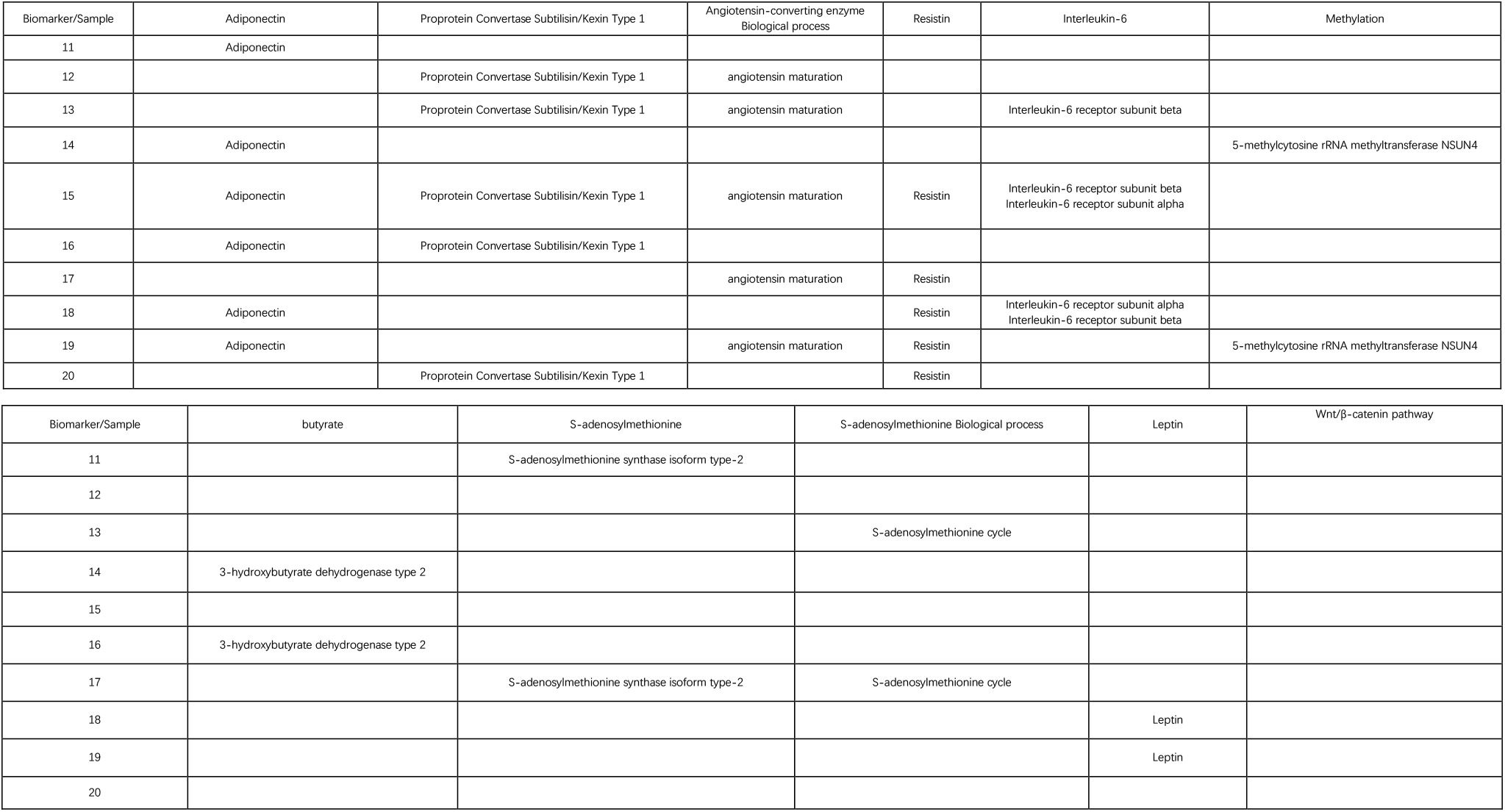
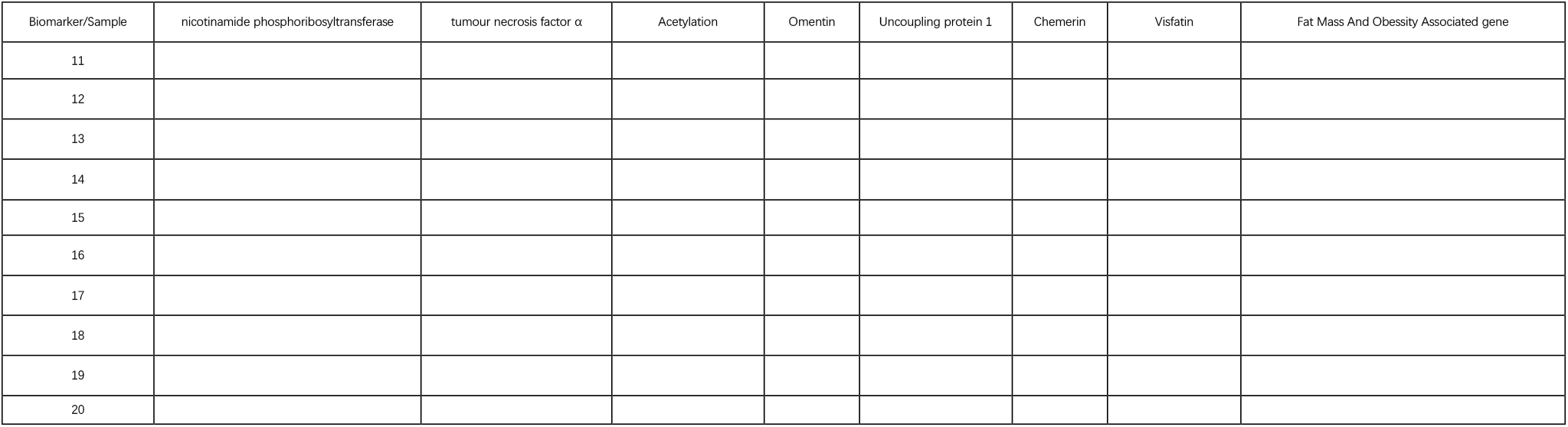
Known markers of obesity possessed by individuals in the obese group.

### 3.4 Discussion

The potential of urine proteome in exploring obesity-related mechanisms, finding potential drug targets and providing personalised treatment options can be seen in this study, and it is worth expanding the sample size further to continue the exploration.

## 4 Conclusion

The urine proteome can distinguish the obese people from the normal weight people, and the differential proteins in the urine proteome have key proteins that are known to be related to obesity and metabolism, and the biological processes of differential proteins also related biological processes such as nutrition and metabolism. Urine proteome has the potential to explore the pathogenesis of obesity and provide personalized treatment.

